# A novel benzoxaborole works by targeting FabI in *Escherichia coli*

**DOI:** 10.1101/2020.12.15.423001

**Authors:** Soma Mandal, Tanya Parish

**Affiliations:** Infectious Disease Research Institute, Seattle, WA, USA; Seattle Children’s Research Institute, Seattle, WA, USA

## Abstract

To combat the looming crisis of antimicrobial-resistant infections, there is an urgent need for novel antimicrobial discovery and drug target identification. The benzoxaborole series were previously identified as inhibitors of mycobacterial growth. Here, we demonstrate that a benzoxaborole is also active against the Gram negative bacterium, *Escherichia coli* in vitro. We isolated resistant mutants of *E. coli* and subjected them to whole genome sequencing. We found mutations in the enoyl acyl carrier protein FabI. Mutations mapped around the active center site located close to the co-factor binding site. This site partially overlaps with the binding pocket of triclosan, a known FabI inhibitor. Similar to triclosan, the interaction of the benzoxaborole with FabI was dependent on the co-factor NAD^+^. Identification of the target of this compound in *E. coli* provides scope for further development and optimization of this series for Gram negative pathogens.

## Introduction

Antimicrobial resistance is a growing problem and at current rates of increase, it is estimated by the year 2050 ∼10 million people will die due to infections caused by resistant bacteria (1). In order to tackle this crisis, we need to identify and characterize novel scaffolds for development as new antimicrobial agents. We previously identified AN11527, a benzoxaborole based compound as a potent inhibitor of *Mycobacterium tuberculosis* growth (2). This is an attractive series, since it has good antibacterial activity against extracellular and intracellular bacteria but lacks cytotoxicity. The target for this series is unknown, although resistant mutants had single nucleotide polymorphisms in three genes (Rv1683 Rv3068c and Rv0047c) (2). Rv0047c is involved in inositol metabolism, Rv3068c encodes a non-essential protein involved in glucose metabolism (PgmA) and Rv1683 encodes a bifunctional protein involved in triglyceride metabolism with both lipase and synthetase activity (3).

In this study we show that AN11527 is active against *Escherichia. coli*. We identified the cellular target of AN11527 in *E. coli* as the enoyl-acyl carrier protein reductase FabI. We determined a potential binding pocket for AN11527 which is located at the active center site and close to the cofactor binding site. We also determined that the binding interactions of AN11527 partially overlaps with that of the NAD^+^-dependent inhibitor Triclosan, but not that of the co-factor independent 4-hydroxy-2-pyridine compound NITD-916.

## Materials and methods

### Determination of antibacterial activity

*Mycobacterium smegmatis* was grown in Middlebrook 7H9 medium supplemented with Middlebrook ADC enrichment (Becton Dickinson), 0.5% v/v glycerol and 0.05% w/v Tween80. *E. coli* was cultured in LB medium. *Staphylococcus aureus* was cultured in MH broth. We determined MICs against *M. smegmatis* mc^2^155, *E. coli* JW5503 and *S. aureus* using a broth microdilution assays. The compound was tested as a two-fold serial dilution at a starting concentration of 100 µM (final DMSO concentration of 2%). Growth was measured by OD and MIC_90_ was recorded at the lowest concentration of compound which inhibited growth by 90% compared to controls.

### Isolation of AN11527-resistant mutants

*E. coli* JW5503 was plated on LB agar containing 4X and 8X MIC AN11527 at 8 x10^8^ to 2×10^9^ CFUs. Single colonies were isolated after overnight incubation at 37°C and cultured in 5 ml LB for genomic DNA isolation. Genomic DNA from resistant isolates was isolated and prepared for sequencing using the Nextera XL library kit. Whole genome sequencing was carried out using 2 × 150 paired-end reads at the Colorado State University whole genome sequencing facility. The *fabI* gene was PCR-amplified using primers 5’-ATGGGTTTTCTTTCCGGTAAGCGCATTC -3’ and 5’-TTATTTCAGTTCGAGTTCGTTCA -3’and sequenced.

### Cloning and purification of *E. coli* FabI

E. *coli FabI* gene was PCR-amplified using primer pair 5’-CATATG ATGGGTTTTCTTTCCGGTAAGCGCAC-3 and 5′-GATCC TTATTTCAGTTCGAGTTCGTTC-3 primers. The PCR product was cloned in to pET15(b) using sites NdeI and BamHI to obtain the plasmid pET15b-FabI expressing FabI with an N-terminal His tag. FabI was expressed in BL21 (DE3) cells as follows: cells were grown to an OD of 0.4 to 0.6, 1 mM IPTG was added and protein was purified from the soluble fraction using a Ni-NTA column with 50 mM HEPES pH 8.0, 500 mM NaCl, 5% glycerol, 1 mM EDTA, 1 mM DTT. The purified protein was concentrated and stored in buffer containing 50mM HEPES pH 8.0, 500 mM NaCl, 50% glycerol, 5 mM DTT. FabI A197G was constructed by site directed mutagenesis using primers (5’ CCGTACTCTGGCGGGATCCGGTATCAAAG 3’) and (5’ CTTTGATACCGGATCCCGCCAGAGTACGG 3’).

### Thermal shift assays

We determined protein melting temperatures using the Prometheus NT.48 instrument (NanoTemper Technologies). Purified FabI was diluted to a concentration of 25 μM in 50 nM HEPES, 200mM NaCl. Compound was added from 1.56 to 200 µM; NADH and NAD^+^ were used at 250 µM. The reaction mixture was incubated at 25 C for 10 mins, 10 μL sample was loaded into the capillaries and placed on the sample holder. A temperature gradient from 20 °C to 80 °C at 0.5 or 1°C per min was applied. Intrinsic protein fluorescence at 330 and 350 nm was recorded and the Tm was calculated using Nano Temper Technologies software.

## Results and Discussion

### The benzoxaborole AN11527 is active against *E. coli*

We determined the activity of a novel benzoxaborole against *M. smegmatis, S. aureus* and *E. coli*. We used a *tolC* mutant strain of *E. coli* (JW5503). Although the compound showed good potency against *M. tuberculosis* (2), it had no activity against the non-pathogenic organism *M. smegmatis* (MIC >200 µM). There was no activity against the Gram positive species, *S. aureus* (MIC >200 µM). However, surprisingly there was good activity against the Gram negative organism, *E. coli* with an MIC of 6.25 µM.

Since *E. coli* was sensitive, we isolated resistant mutants in order to identify the target or mechanism of resistance. We plated 10^8^-10^9^ CFU on agar plates with 25 or 50 µM (4X and 8X MIC) compound and isolated resistant mutants. We confirmed that strains were resistant by streaking onto solid medium with 25 µM AN11527. Resistance was further confirmed in liquid broth and all mutant strains exhibited high-level resistance with > 32-fold shift in MIC (Table 1).

**Table 1.**
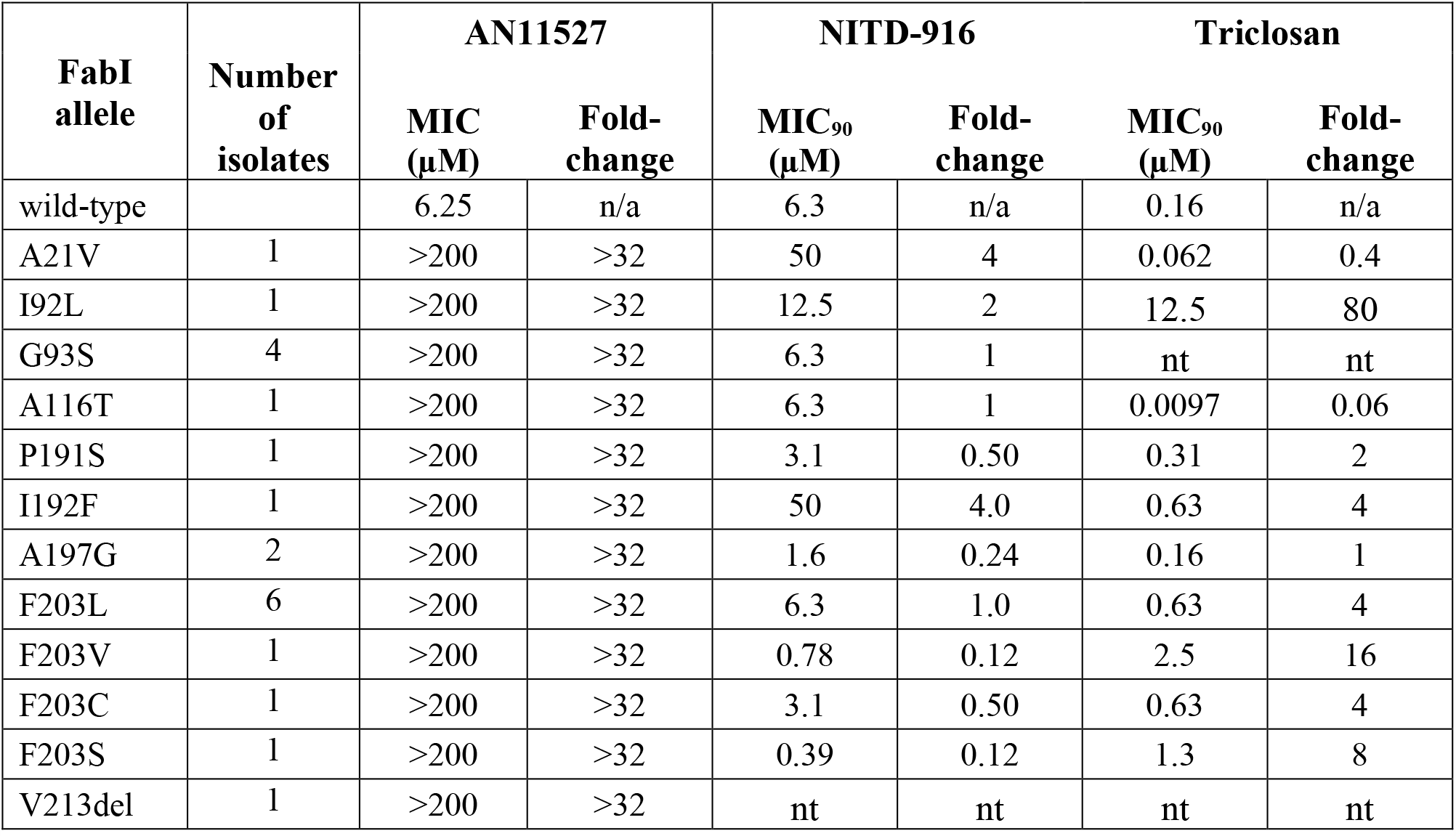
Characterization of resistant mutants. Minimum inhibitory concentrations (MIC) were determined in liquid medium against *E. coli* JW5503 and AN11527-resistant *E. coli* JW5503. Data are from two independent runs.

We conducted whole genome sequencing on two resistant-mutants and the wild-type strain with more than 96% coverage and we identified mutations in *fabI*. Previous work has demonstrated that similar compounds target mycobacterial InhA (4) an ortholog of FabI, and diazaborines are known to inhibit FabI (5). FabI and InhA are involved in the synthesis of fatty acids, although InhA is a enoyl acyl carrier protein reductase involved in the synthesis of mycolic acids which are specific to mycobacteria (6). Thus, it seemed possible that FabI is the target in *E. coli*. We selected 21 AN11527-resistant strains and sequenced *fabI* (Table 1). All of the strains had SNPs in *fabI* covering 9 different amino acid residues with 12 different mutations (11 SNPs and 1 deletion) (Table 1).

The crystal structure of FabI in *E. coli* has been determined (9,12). We mapped the amino acid residues from our resistant isolates onto the three-dimensional structure. The mutations form a cluster large enough to encompass a molecule of AN11527 and are located around the active center site (Figure 1). Several of the residues lie close to the NADH/NAD^+^ binding site (Figure 2) and are involved in the co-factor binding. The binding pocket for AN11527 overlaps with other FabI inhibitors including a 4-hydroxy-2-pyridine (NITD-916), isoniazid, and triclosan (7–9) (Figure 3). Therefore, we determined whether the mutations also conferred resistance to triclosan. We observed a range of resistance for the different mutations, with SNPs in I92, I192, F203 all conferring >4-fold resistance (Table 1). The I92L allele conferred high level resistance (>80-fold), whereas mutations in F203 had varying resistance from 4-16-fold. Interestingly, the A116T allele conferred susceptibility to triclosan.

**Figure 1.**
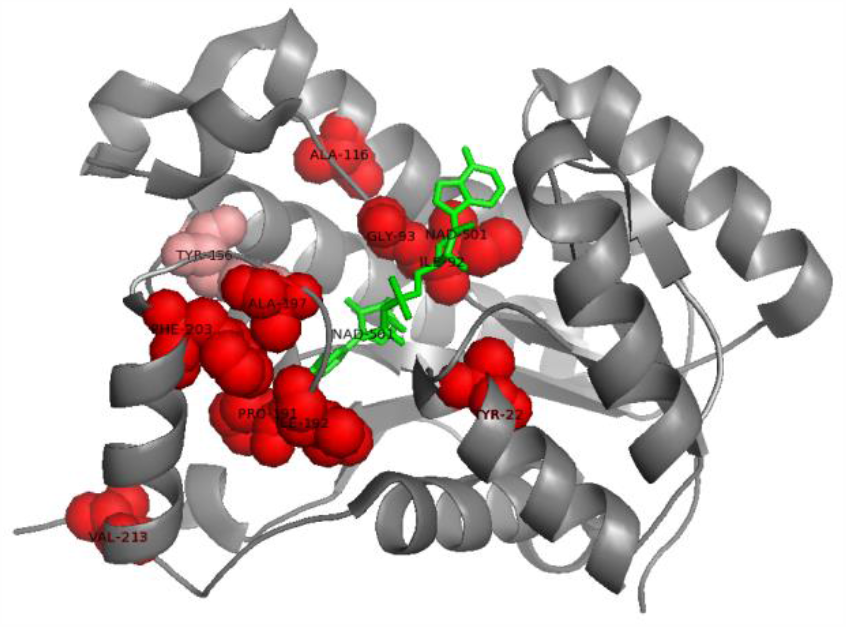
Crystal structure of FabI (1D8A). (A) Residues that confer resistance to AN11527 are shown in red and NAD^+^ in green. The catalytic residue Y156 is in pink. Residues that confer resistance to AN11527 are highlighted in red and are located around the catalytic residue, the active center and close to the NAD+ binding site.

**Figure 2:**
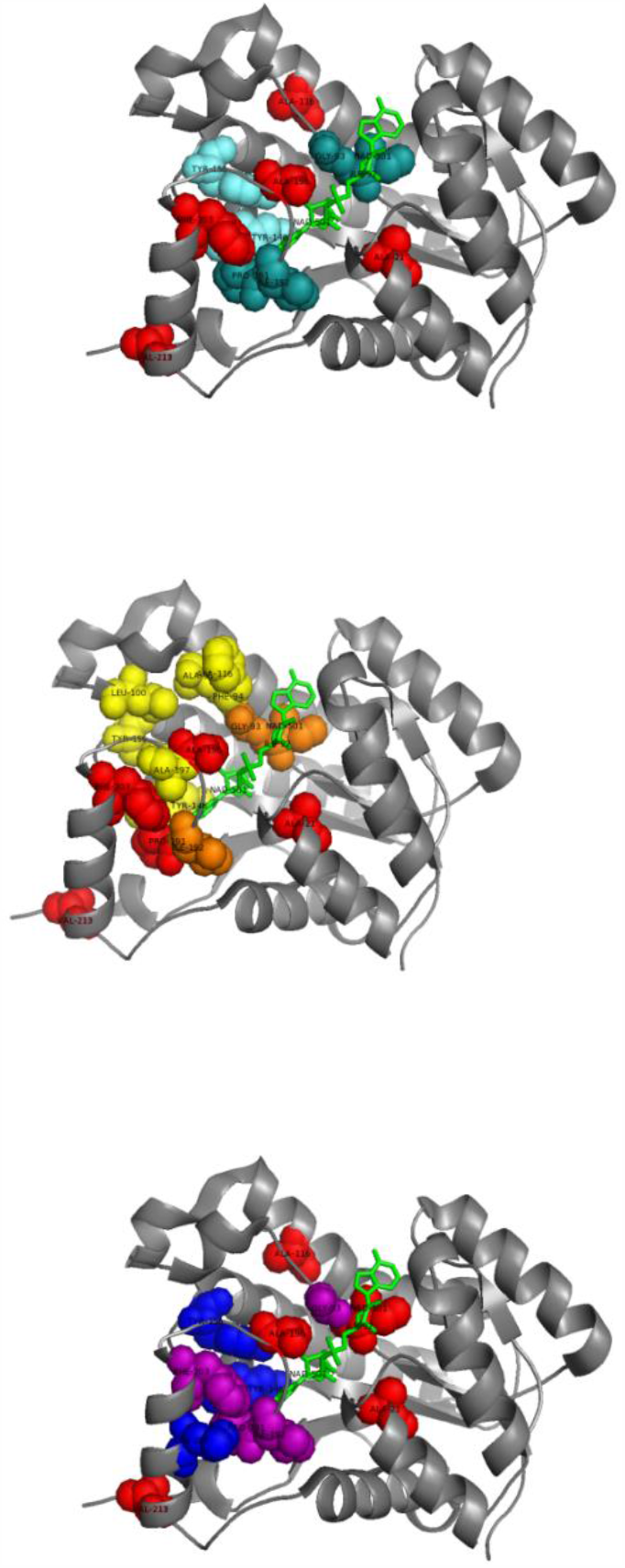
Comparison of the binding pockets for FabI inhibitors. The proposed binding pocket of each inhibitor occupies the same general area around the active center site and is close to the NAD+ binding location (FabI crystal structure 1D8A). Residues that confer resistance to AN11527 are in red. NAD^+^ is shown bound. For isoniazid and NITD-916 interacting residues are selected based on FabI and InhA homology. (A) Residues that confer resistance to isoniazid are light green; residues that confer resistance to both isoniazid and AN11527 are in dark green. (B) Residues that confer resistance to triclosan are light yellow; residues that confers resistance to both isoniazid and AN11527 are dark orange. (C) Residues that confer resistance to NITD-916 are blue; residues that confer resistance to both NITD-916 and AN11527 are purple.

**Figure 3:**
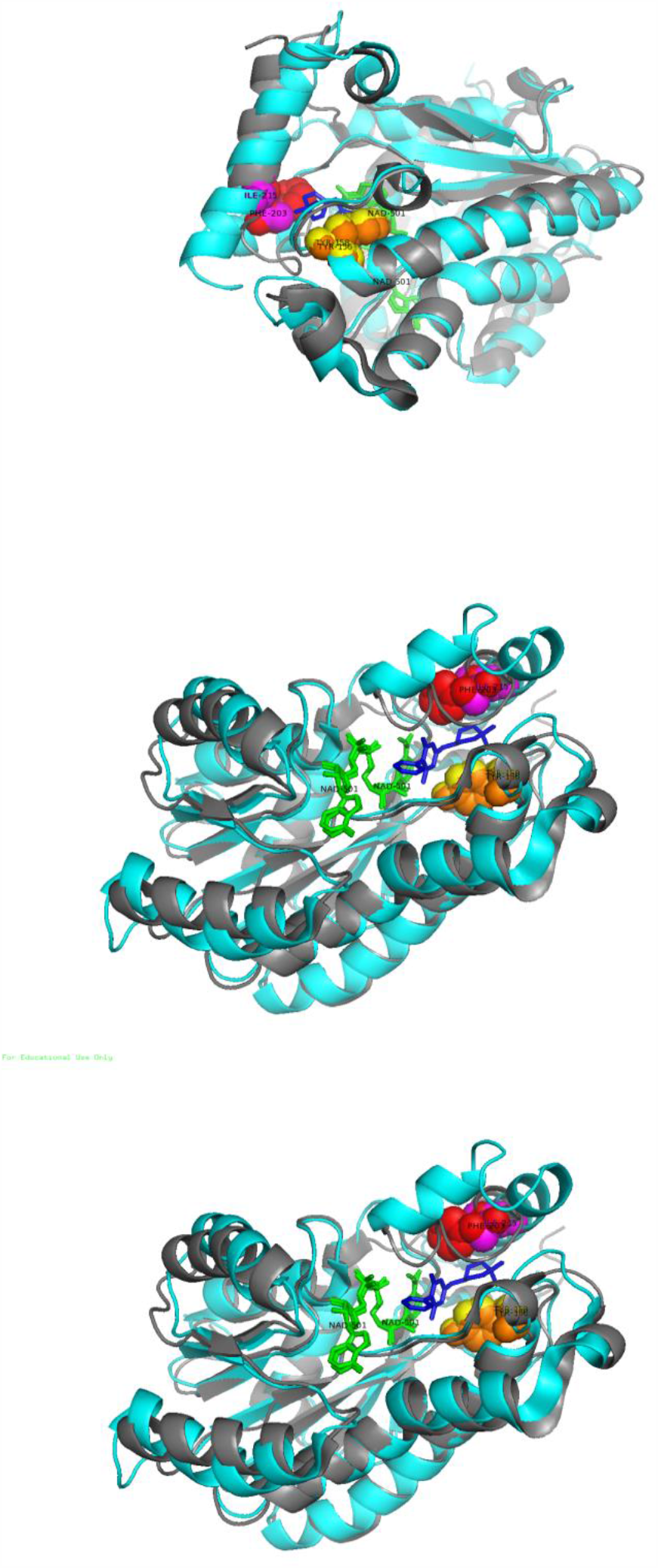
Location of FabI F203 and InhA I215. Superimposition of FabI and InhA structures; *E. coli* FabI with triclosan in grey and *M. tuberculosis* InhA with NITD-916 in cyan. In yellow and orange are the catalytic residue Y156 and in red and purple are the residue FabI F203 and InhA I215 respectively.

### Mycobacterial InhA inhibitors are active against *E. coli*

Previous work suggested that the 4-hydroxy-2-pyridine compound NITD-916 is active against *M. tuberculosis*, but not *E. coli* (7). Since the benzoxaborole was active against *E. coli* and there is good structural similarity between FabI and InhA, we tested it for activity against the TolC mutant strain. We found that NITD-916 had good activity (MIC = 6.25 µM; Table 1) and thus its lack of activity is likely due to efflux and lack of access to the target, as has been demonstrated for other FabI inhibitors in *E. coli* (10). We tested our mutant isolates for cross-resistance to NITD-916; only two mutants showed low level resistance (A21V and I192F) (Table 1). Both of these residues interact with the co-factor and this may have an indirect effect on AN11527 and NITD-916 binding. However, several mutants were hypersensitive (A197G, F203V and F203S). The equivalent of FabI F203 in M. *tuberculosis* is I215 which is involved in NITD-916 binding. However, in *E. coli* the InhA_F203_ mutant strain did not exhibit cross-resistance or hypersensitivity to NITD-916 (Figure 4). Thus, even though the binding pocket of NITD-916 and AN11527 overlaps, the lack of cross-resistance suggests differences in binding.

### AN11527 binding to FabI is co-factor dependent

FabI is a fatty acid reductase which uses cellular NADH as a co-factor to reduce the enoyl carbon of fatty acids (8). FabI inhibitors can be divided in two groups based on whether their binding is co-factor dependent or independent. Depending upon the mechanism of interaction bacterial FabI inhibitors are categorized in two groups, one that requires a co-factor (NAD^+^/NADH) to bind and another class which binds directly (11).

We determined whether benzoxaborole binding was co-factor dependent using thermal shift assays. NAD^+^-dependent binding of triclosan resulted in a large thermal shift (Table 2). No changes in T_m_ were observed when incubated with AN11527 alone or AN11527 plus NADH. However, AN11527 in the presence of NAD^+^ resulted in a shift. We confirmed this shift was dose-dependent (Table 3). We also tested the FabI_A197G_ allele for binding. As expected, no thermal shift was seen in the presence of AN11527 even in the presence of NAD^+^, confirming loss of binding. Triclosan was still able to bind to the mutant protein in the presence of NAD^+^ as expected from the lack of resistance of the strain. These data further support that FabI is the target of AN11527 in *E. coli*.

**Table 2.** Interaction of FabI with AN11527. FabI melting temperatures were measured by nanoDSF. Each sample contained 25 nM FabI plus compound at 200 µM and NAD+ at 250 µM. where stated. The inflection point in °C was recorded as the melting temperature; shifts are the difference in Tm. Data are representative of two independent experiments.

| Allele | Molecule | Concentration | NAD<sup>+</sup> | T<sub>m</sub> | Shift |
| --- | --- | --- | --- | --- | --- |
| **FabI<sub>wt</sub>** | None |  | - | 52.1 |  |
|  | AN11527 |  | - | 52.1 |  |
|  | None |  | + | 52.1 |  |
|  | AN11527 |  | + | 56 | 3.9 |
|  | Triclosan |  | + | 71.9 | 19.8 |
| **FabI<sub>A197G</sub>** | None |  | - | 51.4 |  |
|  | AN11527 |  | - | 51.2 |  |
|  | None |  | + | 51.3 |  |
|  | AN11527 |  | + | 52.2 | 0.9 |
|  | Triclosan |  | + | 65 | 13.7 |

**Table 3.**
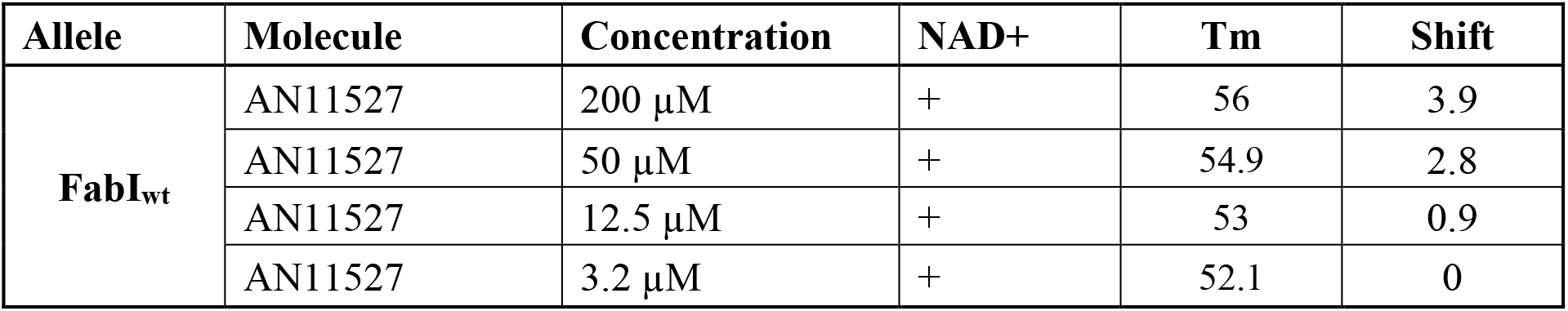
Dose-dependent thermal shift of FabI with AN11527. FabI melting temperatures were measured by nanoDSF. Each sample contained 25 µM FabI plus compound and 250 µM NAD^+^. The inflection point in °C was recorded as the melting temperature; shifts are the difference in Tm. Data are representative of two independent experiments.

| Allele | Molecule | Concentration | NAD<sup>+</sup> | T<sub>m</sub> | Shift |
| --- | --- | --- | --- | --- | --- |
| **FabI<sub>wt</sub>** | AN11527 | 200 μM | + | 56 | 3.9 |
|  | AN11527 | 50 μM | + | 54.9 | 2.8 |
|  | AN11527 | 12.5 μM | + | 53 | 0.9 |
|  | AN11527 | 3.2 μM | + | 52.1 | 0 |

## Conclusion

FabI is an essential enzyme in many bacterial species, as it catalyzes a key step in the biosynthesis of fatty acid pathway as part of the FAS II complex. It is interesting to note that although the substrates and products for FabI and the mycobacterial ortholog InhA are different, a number of inhibitors are active against both enzymes (11). A wide range of FabI/InhA inhibitors have been identified including the frontline tuberculosis drug isoniazid and the antibacterial agent triclosan, as well as series under development such as the 4-hydroxy-2-pyridine compound NITD-916. We demonstrate that a benzoxaborole compound and NITD-916 are both active against the *E. coli* target and that resistance maps to FabI. We demonstrate that AN11527 binds to InhA in a co-factor dependent manner which is abrogated in mutant proteins.

## References

1. O’Neil J. 2014. Review on antibiotic resisitance. Antimicrobial Resistance?: Tackling a crisis for the health and wealth of nationsHealth and Wealth Nations.

2. Patel N, O’Malley T, Zhang YK, Xia Y, Sunde B, Flint L, Korkegian A, Ioerger TR, Sacchettini J, Alley MRK, Parish T. 2017. A novel 6-benzyl ether benzoxaborole is active against mycobacterium tuberculosis in vitro. Antimicrob Agents Chemother 61.

3. Low KL, Shui G, Natter K, Yeo WK, Kohlwein SD, Dick T, Rao SPS, Wenk MR. 2010. Lipid droplet-associated proteins are involved in the biosynthesis and hydrolysis of triacylglycerol in Mycobacterium bovis bacillus Calmette-Guérin. J Biol Chem 285:21662–21670.

4. Xia Y, Zhou Y, Carter DS, McNeil MB, Choi W, Halladay J, Berry PW, Mao W, Hernandez V, O’Malley T, Korkegian A, Sunde B, Flint L, Woolhiser LK, Scherman MS, Gruppo V, Hastings C, Robertson GT, Ioerger TR, Sacchettini J, Tonge PJ, Lenaerts AJ, Parish T, Alley MRK. 2018. Discovery of a cofactor-independent inhibitor of Mycobacterium tuberculosis InhA. Life Sci Alliance 1:e201800025.

5. Jordan CA, Sandoval BA, Serobyan M V., Gilling DH, Groziak MP, Xu HH, Vey JL. 2015. Crystallographic insights into the structure-activity relationships of diazaborine enoyl-ACP reductase inhibitors. Acta Crystallogr Sect Struct Biol Commun 71:1521–1530.

6. Dessen A, Quémard A, Blanchard JS, Jacobs WR, Sacchettini JC. 1995. Crystal structure and function of the isoniazid target of Mycobacterium tuberculosis. Science (80-) 267:1638–1641.

7. Manjunatha UH, Rao SPS, Kondreddi RR, Noble CG, Camacho LR, Tan BH, Ng SH, Ng PS, Ma NL, Lakshminarayana SB, Herve M, Barnes SW, Yu W, Kuhen K, Blasco F, Beer D, Walker JR, Tonge PJ, Glynne R, Smith PW, Diagana TT. 2015. Direct inhibitors of InhA are active against Mycobacterium tuberculosis. Sci Transl Med 7:269ra3.

8. Rafi S, Novichenok P, Kolappan S, Zhang X, Stratton CF, Rawat R, Kisker C, Simmerling C, Tonge PJ. 2006. Structure of acyl carrier protein bound to FabI, the FASII enoyl reductase from Escherichia coli. J Biol Chem 281:39285–39293.

9. Sivaraman S, Zwahlen J, Bell AF, Hedstrom L, Tonge PJ. 2003. Structure-activity studies of the inhibition of FabI, the enoyl reductase from Escherichia coli, by Triclosan: Kinetic analysis of mutant FabIs. Biochemistry 42:4406–4413.

10. Parker EN, Drown BS, Geddes EJ, Lee HY, Ismail N, Lau GW, Hergenrother PJ. 2020. Implementation of permeation rules leads to a FabI inhibitor with activity against Gram-negative pathogens. Nat Microbiol 5:67–75.

11. Lu H, Tonge PJ. 2008. Inhibitors of FabI, an enzyme drug target in the bacterial fatty acid biosynthesis pathway. Acc Chem Res 41:11–20.

12. Qiu X, Janson CA, Court RI, Smyth MG, Payne DJ, Abdel-Meguid SS. Molecular basis for triclosan activity involves a flipping loop in the active site. Protein Sci. 1999 Nov;8(11):2529–32. doi: 10.1110/ps.8.11.2529. PMID: 10595560; PMCID: PMC2144207.

